# Structural basis of nirmatrelvir and ensitrelvir resistance profiles against SARS-CoV-2 Main Protease naturally occurring polymorphisms

**DOI:** 10.1101/2022.08.31.506107

**Authors:** Gabriela Dias Noske, Ellen de Souza Silva, Mariana Ortiz de Godoy, Isabela Dolci, Rafaela Sachetto Fernandes, Rafael Victório Carvalho Guido, Peter Sjö, Glaucius Oliva, Andre Schutzer Godoy

## Abstract

SARS-CoV-2 is the causative agent of COVID-19. M^pro^ is the main viral protease, with a critical role in replication and, therefore, an attractive target for antiviral drug discovery. The clinically approved drug nirmatrelvir from Pfizer, and the clinical candidate ensitrelvir from Shionogi Pharmaceuticals had so far showed great potential for treatment of viral infections. Despite the importance of new therapeutics, the broad use of antivirals is often associated with mutation selection and resistance generation. Herein, we characterized 14 naturally occurring polymorphisms that are already in circulation and are within the radius of action of these two antivirals. Nirmatrelvir retained most of its *in vitro* activity against most polymorphism tested, while mutants G143S and Q189K were associated with higher resistance. For ensitrelvir, higher resistance was observed for polymorphisms M49I, G143S and R188S, but not for Q189K, suggesting a distinct resistance profile difference between the two inhibitors. The crystal structures of selected polymorphism reveal the structural basis for resistance generation. Our data will assist the monitoring of potential resistant strains, support the design of combined therapy to avoid resistance, as well as assist the development of a next generation of M^pro^ inhibitors

## 1. Introduction

SARS-CoV-2 is a highly transmissible β-coronavirus (1, 2) with a genome composed by a single RNA positive strand that comprises about 30 kb encoding for 16 nonstructural, 4 structural and 6 accessory proteins(3). The viral replicase codifies two frame shifting open reading frames, ORF1a/ORF1ab containing 16 non-structural proteins required for viral replication(4). The SARS-CoV-2 Main protease (M^pro^) or 3C-like protease (3CL^pro^) is a dimeric cysteine protease responsible for the cleavage of the viral polyproteins 1a and 1ab in 11 sites, including its own N and C-terminal (5–7). M^pro^ substrate recognition has unique features and is specific for Gln residue at P1, hydrophobic residues at P2 and small side chains such as Ser, Ala, Gly, Leu at P1’(8). The absence of similar sites in human proteases together with the importance of the enzyme for the viral replication makes M^pro^ a primary target for antiviral discovery and development(9).

Several small molecules were identified as M^pro^ inhibitors that exhibited efficacy in cellular culture including boceprevir, carmofur, MAT-POS-e194df51-1, PF-07321332 (nirmatrelvir) and S-217622 (ensitrelvir)(6, 9–14). The first oral COVID-19 antiviral from Pfizer, Paxlovid^™^, is a combination of nirmatrelvir and ritonavir, a CYP3A4 inhibitor, with safety, and efficacy demonstrated in clinical trials, and approved for use by FDA in December 2021(9, 15). In addition, the compound ensitrelvir from Shionogi, is a promising non-covalent inhibitor of M^pro^ (12). Currently in phase 3 of clinical trials, the compound has showed exciting pharmacokinetics properties, with potential for therapeutical doses to be reached without requirement of CYP inhibitors (12).

Like other viruses, SARS-CoV-2 genome is constantly mutating during replication. Although most mutations are not expected to generate a viable variant(16), WHO is constantly monitoring the emergence of SARS-CoV-2 mutations, since recent variants had exhibited more transmissible and infectious properties, and can affect the vaccines effectiveness(17, 18). Moreover, amino acid replacements in the viral targets, can impact the catalytic activity of the enzyme and modify the efficacy of inhibitors(19, 20). The *in vitro* effectiveness of nirmatrelvir against variants Alpha (B.1.1.7), Beta (B.1.351), Delta (B1.617.2), Gamma (P.1), Lambda (B.1.1.1.37/C37), Omicron (B.1.1.529) has already been demonstrated (21). However, none these variants contain mutations that are within the radius the active site. Herein, we evaluated the effect of active site single mutations from circulating polymorphisms SARS-CoV-2 M^pro^ on the enzyme kinetic and nirmatrelvir/ensitrelvir efficacy. We also used X-ray crystallography to characterize the structural features of selected polymorphism. These findings provided key information for predicting and avoiding resistance, designing the next generation of inhibitors, and raised important considerations for combination therapies.

## 2. Results

### 2.1. Identification of circulating SARS-CoV-2 M^pro^ active site polymorphisms

For this study, we selected active site polymorphic versions of M^pro^ that have been already identified in circulation. In this sense, we assessed sequencing data available from GISAID hCoV-19/SARS-CoV-2 sequences database(22) (containing approximately 7 million genomes as of December 20th, 2021) in CoV-GLUE (http://cov-glue.cvr.gla.ac.uk) relative to M^pro^. We identified 389 distinct polymorphisms in M^pro^ with a n ≥ 10 individuals. From those, and based on available structural information (13), we selected mutants that were within 8.0 Å of radius from each inhibitor. We identified 16 polymorphisms, which are summarized in Figure 1. Among these, four key nirmatrelvir contacts residues were identified (e.g., M49, G143, M165 and Q189). To investigate the impacts of the polymorphisms on M^pro^ activity and inhibitors efficacy, we were able to characterize 15 mutants. Mutant N142L did not generated any soluble protein, and therefore not characterized further.

**Figure 1.**
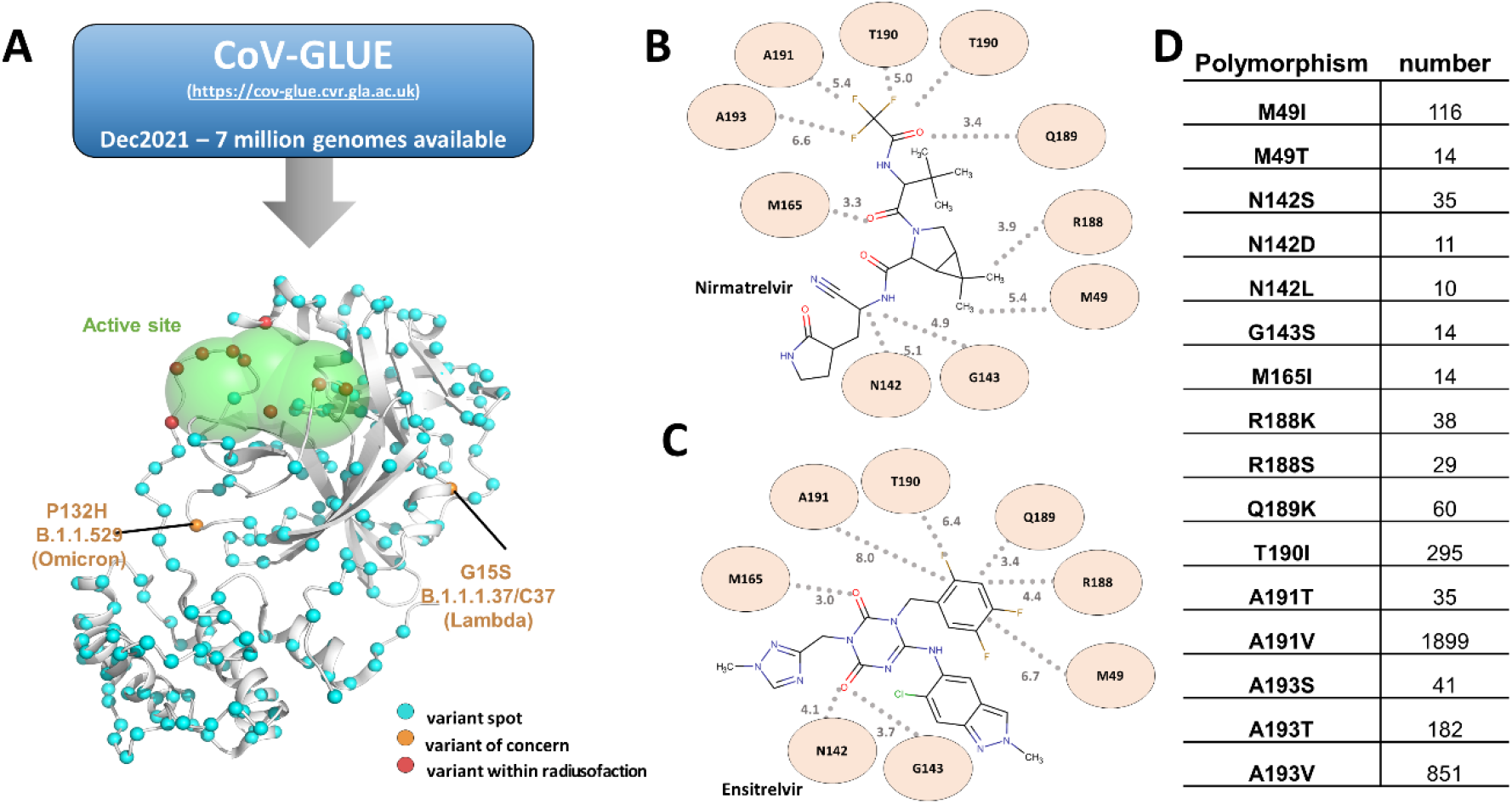
SARS-CoV-2 M^pro^ amino acids polymorphisms identified in the genomic database. (A) M^pro^ model showing variant spots identified in the genomic database, using a threshold of n ≥ 10. The Cα of identified variants spots are showed as cyan spheres, while red spheres are those variants within radius of 8 Å ligands. Orange spheres show spots for variants of concern. Active site of M^pro^ is colored with a green blob. (B) 2D plot of nirmatrelvir showing the distance and closest contacts with selected polymorphisms. (B) 2D plot of ensitrelvir showing the distance and closest contacts with selected polymorphisms. (D) Panel containing polymorphism name, and substitution, and number of individuals identified (n).

### 2.2. Enzymatic characterization of SARS-CoV-2 M^pro^ polymorphisms

M^pro^ mutants were active and able to recognize and cleave the fluorogenic substrate. The assessed kinetic parameters of all mutants (*K*_M_, *k*_cat_ and the relative efficiency) are summarized in Table 1 and the Michaelis-Menten plots are shown in Fig. S1. The WT enzyme showed a *K*_M_ value of 22 ± 2 (µM), and a *k*_cat_ value of 31 ± 1, which agrees with previous characterizations of M^pro^ for this substrate(7). The mutant panel exhibited *K*_M_ values ranging from 6.4 – 25.4 µM, and relative efficiencies (*K*_M_/*k*_cat_) relative to the WT that ranged between 3 – 205 % (Table 1). Seven mutants (M49I, M49T, N142D, M165I, R188K, T190I and A191T) exhibited a catalytic efficiency significantly greater than wild type M^pro^, with relative efficiencies increments between 1 to 2-fold, where N142D and R188K mutants were the ones with greater relative efficiencies. The other five mutants (N142S, R188S, A191V, A193S and A193T) showed similar catalytic efficiency to the wild type (Fig. 2A). Mutants G143S, Q189K and A193V showed decreased catalytic efficiencies in comparison with the wild type, exhibiting activity reduced by 33, 3 and 2-fold, respectively (Fig. 2A).

**Table 1.** Kinetic parameters of M^pro^ mutants. Relative efficiency is the k_cat_/*K*_*m*_ of polymorphisms relative to wild type M^pro^.

| Polymorphism | $K_m$ (μM) | $k_{cat}$ (RFU/μM.s) | Relative Efficiency |
| --- | --- | --- | --- |
| M49I | 6 ± 1 | 16.0 ± 0.6 | 1.8 ± 0.2 |
| M49T | 7 ± 2 | 19 ± 1 | 1.8 ± 0.2 |
| N142S | 15 ± 3 | 25 ± 1 | 1.2 ± 0.2 |
| N142D | 9 ± 1 | 27 ± 2 | 2.0 ± 0.1 |
| G143S | 25 ± 11 | 1.0 ± 0.1 | 0.030 ± 0.001 |
| M165I | 7.7 ± 0.7 | 19.6 ± 0.5 | 1.8 ± 0.1 |
| R188K | 8.2 ± 0.9 | 23 ± 1 | 2.00 ± 0.08 |
| R188S | 22 ± 4 | 28 ± 2 | 0.9 ± 0.3 |
| Q189K | 9 ± 1 | 4.5 ± 0.2 | 0.30 ± 0.09 |
| T190I | 10 ± 2 | 23 ± 1 | 1.7 ± 0.2 |
| A191T | 11 ± 1 | 25 ± 1 | 1.7 ± 0.2 |
| A191V | 18 ± 3 | 29 ± 2 | 1.3 ± 0.3 |
| A193S | 14 ± 2 | 21.5 ± 0.8 | 1.1 ± 0.2 |
| A193T | 14 ± 2 | 22.1 ± 0.9 | 1.0 ± 0.2 |
| A193V | 13 ± 2 | 7.5 ± 0.3 | 0.4 ± 0.1 |
| Wild Type | 22 ± 2 | 30.8 ± 0.8 | 1.0 ± 0.2 |

**Figure 2.**
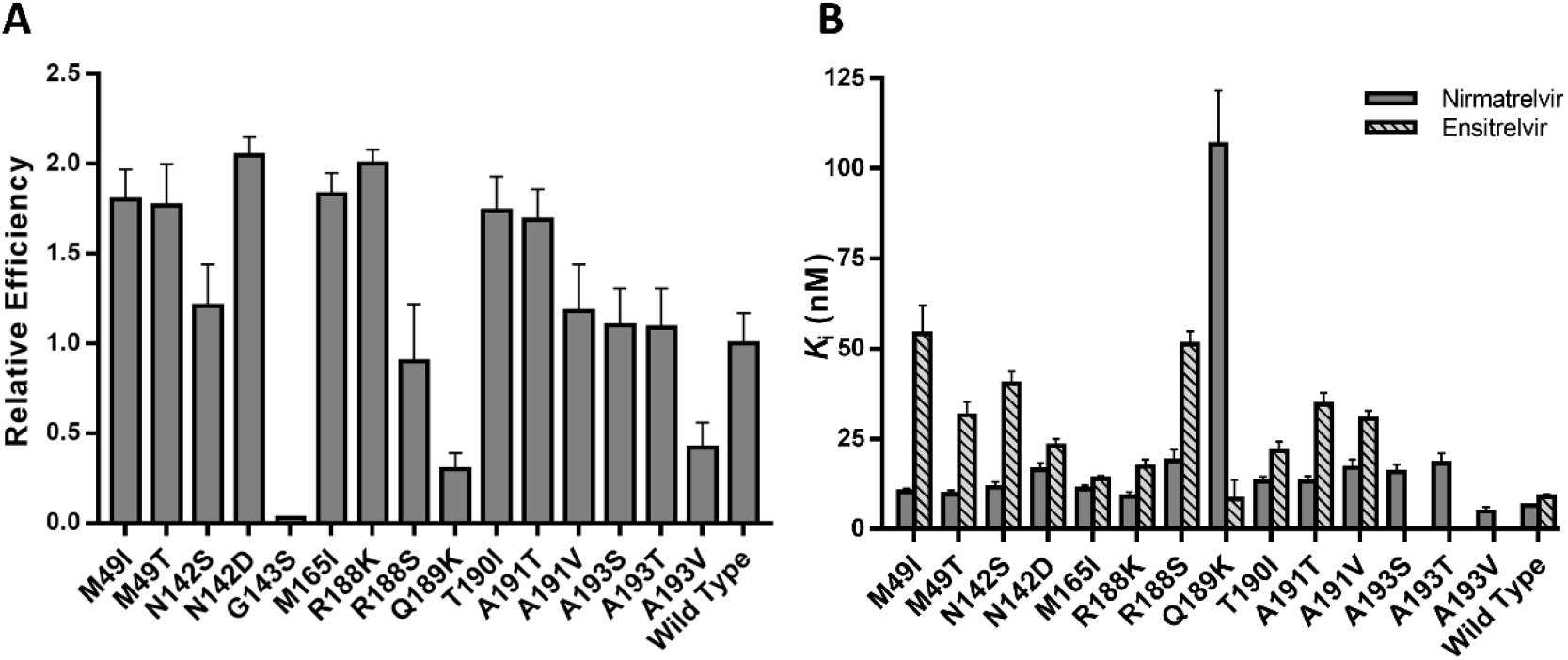
In-vitro characterization of SARS-CoV-2 M^pro^ mutants. (A) Relative catalytic efficiencies of M^pro^ mutants against FRET substrate. Error bars indicate the standard deviation of the triplicates. (B) *K*_i_ determination of nirmatrelvir and ensitrelvir against M^pro^ mutants. Values were calculated using the results obtained in the FRET-based activity assay. For a better visualization, *K*_i_ values of mutant G143S are not represented in this graph

### 2.3. In vitro inhibitory activity of nirmatrelvir against M^pro^ polymorphisms

Next, to access the potential impact of the selected mutations on nirmatrelvir potency, we evaluated the inhibitory activity against M^pro^ polymorphisms through a FRET-based activity assay. Our assay confirmed that nirmatrelvir is a nanomolar inhibitor of M^pro^, with an IC_50_ value of 0.022 ± 0.004 µM and calculated *K*_i_ of 0.006 µM against the WT enzyme. This finding is in good agreement with the reported nirmatrelvir inhibitory activity (9).

The inhibition constant (*K*_i_) observed for most of the mutants, allowed to conclude that nirmatrelvir inhibited the enzymatic activity of all tested polymorphic entities, with *K*_i_ values ranging from 0.005 – 0.106 µM. The only exception was the G143S mutant, where nirmatrelvir *K*_i_ value was 0.96 µM. Mutants M49I, M49T, N142S, M165I, R188K, T190I, A191T and A193V showed higher susceptibility to nirmatrelvir, with *K*_i_ values ranging from 0.005 – 0.013 µM, and no more than 2-fold decrease in the *K*_i_ values relative to the WT was observed (Fig. 2B). N142D, R188S, A191V, A193S and A193T mutants showed modest resistance against nirmatrelvir, with 2 to 3-fold decrease in *K*_i_ values when compared with the WT. Mutants Q189K and G143S exhibited the highest resistance profiles against nirmatrelvir, showing 16 and 147-fold increase in the *K*_i_ values, respectively. The results of the inhibitory effect of nirmatrelvir are summarized in Table 2.

**Table 2.** Inhibition of nirmatrelvir and ensitrelvir against SARS-CoV-2 M^pro^ polymorphisms. IC_50_ Fold Increase is relative to the WT.

| Polymorphism | Nirmatrelvir |  |  | Ensitrelvir |  |  |
| --- | --- | --- | --- | --- | --- | --- |
| | IC<sub>50</sub> (nM) | IC<sub>50</sub> Fold Increase | $K_i$ (nM) | IC<sub>50</sub> (nM) | IC<sub>50</sub> Fold Increase | $K_i$ (nM) |
| M49I | 17.1 ± 0.5 | 0.8 | 10.4 | 139 ± 8 | 10.7 | 54.2 |
| M49T | 16.9 ± 0.5 | 0.8 | 9.6 | 73.0 ± 0.3 | 5.6 | 31.6 |
| N142S | 29 ± 1 | 1.3 | 11.5 | 67 ± 2 | 5.1 | 40.4 |
| N142D | 32 ± 1 | 1.4 | 16.45 | 48.0 ± 0.2 | 3.7 | 23.2 |
| G143S | 3380 ± 128 | 154.4 | 960.2 | 197 ± 0.5 | 15.1 | 141.3 |
| M165I | 19 ± 1 | 0.9 | 11.1 | 32.0 ± 0.4 | 2.5 | 13.9 |
| R188K | 17 ± 1 | 0.7 | 9.0 | 38 ± 3 | 2.9 | 17.2 |
| R188S | 61 ± 3 | 2.8 | 18.95 | 74 ± 1 | 5.7 | 51.3 |
| Q189K | 203 ± 16 | 9.3 | 106.7 | 38 ± 9 | 2.9 | 8.3 |
| T190I | 25 ± 1 | 1.4 | 13.2 | 44 ± 2 | 3.4 | 21.6 |
| A191T | 27 ± 1 | 1.2 | 13.2 | 67 ± 2 | 5.1 | 34.6 |
| A191V | 46 ± 2 | 2.1 | 16.9 | 48.0 ± 0.1 | 3.7 | 30.7 |
| A193S | 27 ± 1 | 1.2 | 15.8 | N.D. | N.D. | N.D. |
| A193T | 32 ± 1 | 1.5 | 18.2 | N.D. | N.D. | N.D. |
| A193V | 11 ± 2 | 0.5 | 4.8 | N.D. | N.D. | N.D. |
| Wild Type | 21.9 ± 0.4 | 1.00 | 6.5 | 13 ± 1 | 1.00 | 8.9 |

N.D.: Not determined

### 2.3. In vitro inhibitory activity of ensitrelvir against M^pro^ polymorphisms

We also assessed the ensitrelvir inhibitory activity against M^pro^ polymorphisms through a FRET-based activity assay. Our assay confirmed ensitrelvir as a potent nanomolar inhibitor of M^pro^, with IC_50_ value of 0.013 ± 0.004 µM and calculated *K*_i_ of 0.009 µM against the WT enzyme, similar to previously described (13).

Ensitrelvir also retained its inhibitory effects against our mutant in the panel, with *K*_i_ values ranging from 0.009 – 0.54 µM. Notably, the exception of the G143S mutant, again, with a *K*_i_ value of 0.141 µM (Fig 2B). None of the mutants exhibited a significant higher susceptibility to ensitrelvir, except for Q189K, with a *K*_i_ value of 0.008 µM, which is comparable with the *K*_i_ determined for the WT. N142D, R188K and T190I mutants showed modest resistance against ensitrelvir, with 2 to 3-fold decrease in *K*_i_ when compared with the WT. M49I, G143S and R188S mutants showed the greatest resistance profiles against ensitrelvir, showing 6, 15, and 6-fold increase in the *K*_i_ values, respectively. The results of inhibitory effect of ensitrelvir are summarized at Table 2.

### 2.4. Crystal structures of mutants in complex with nirmatrelvir and ensitrelvir

For an in-depth understanding of structural basis involved in the resistance profile of the M^pro^ mutants toward nirmatrelvir and ensitrelvir, we determined the crystal structure of selected M^pro^ mutants in complex with both drugs. For nirmatrelvir complexes, we determined the crystals structures of the WT enzyme, as well as M49I, N142S, G143S, Q189K, A193T and A193S mutants at 2.1, 1.9, 1.8, 1.6, 2.4. 2.5 and 1.9 Å resolution, respectively. All crystal structures were obtained in the orthorombic space group with one dimer of M^pro^ in the asymmetric unit, following the pattern of the seeding samples used, whereas the G143S mutant was solved in monoclinic space group with a similar crystal packing, but with two dimers in the asymmetric unit. The structures were refined to *R*_work_/*R*_free_ values of 0.19/0.23, 0.20/0.24, 0.23/0.26, 0.21/0.25, 0.22/0.30, 0.22/0.27 and 0.20/0.24, respectively. Data collection and refinement statistics for the mutants in complex with nirmatrelvir are summarized in Table S2. For ensitrelvir, we determined the crystal complexes of WT and M49I mutant at 2.3 and 2.0 Å and refine those to *R*_work_/*R*_free_ 0.22/0.27 and 0.22/0.25, respectively. Data collection and refinement statistics for WT and M49I mutant in complex with ensitrelvir are summarized in Table S3.

In general, nirmatrelvir exhibited a similar binding mode in all mutants, maintaining most of the key interactions with M^pro^ binding site residues. The electron density around C145 clearly indicated the presence of a covalent bond between the nitrile carbon of nirmatrelvir and the Sγ atom of C145 (1.8 Å, Fig. 3). The WT structure in complex with nirmatrelvir shows key interactions between protein and ligand, including three polar contacts between the pyrrolidone group and amino acids F140 (3.3 Å), H163 (2.6 Å) and E166 (3.1 Å), as well as other two hydrogen bonds between E166 N (2.9 Å) and O (2.8 Å) atoms and the tert-butyl moiety, and a salt bridge between Nirmatrelvir O atom and Q189 from the carbonyl group (4.5 Å) (Fig. 3).

**Figure 3.**
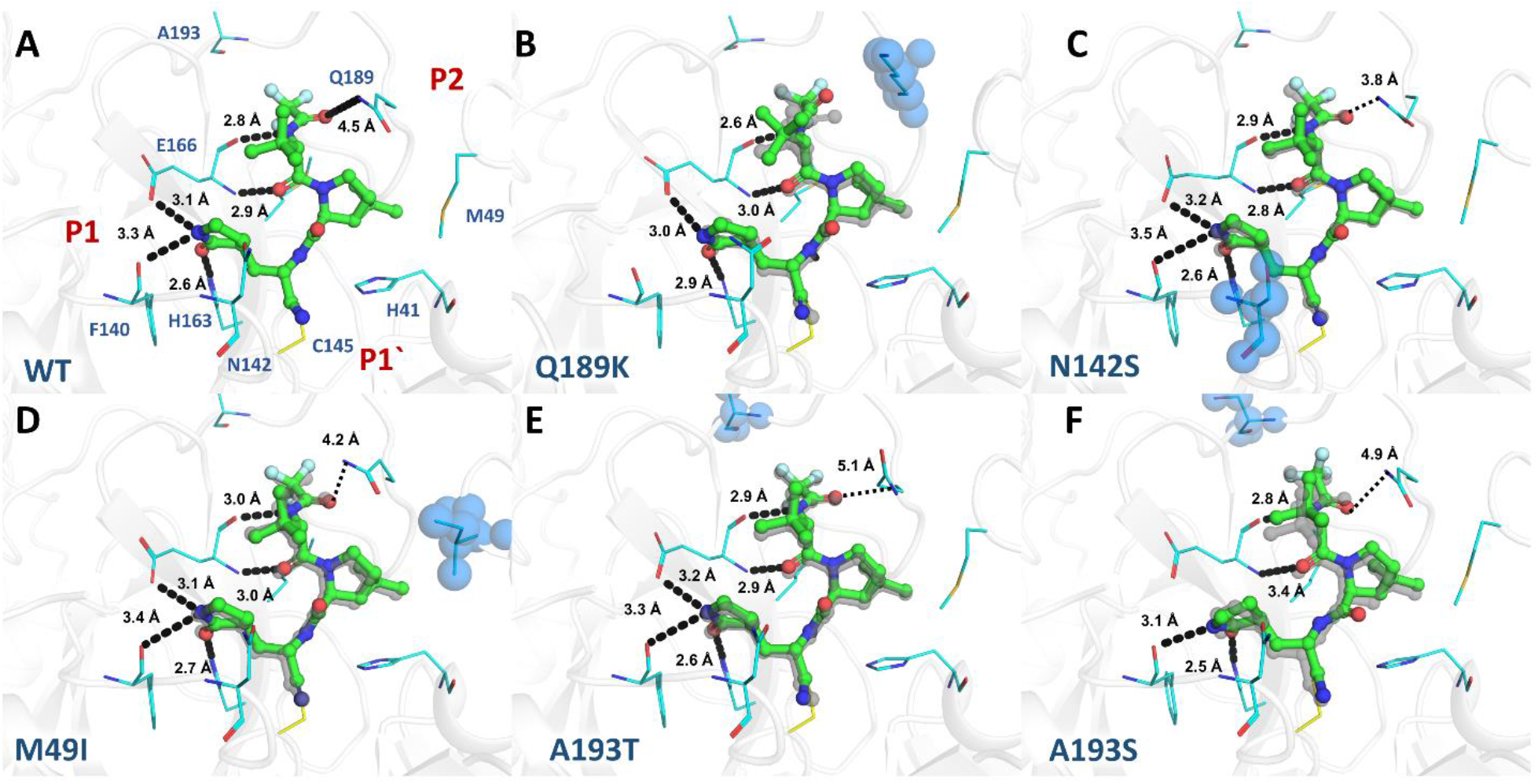
Crystal structures of M^pro^ mutants in complex with nirmatrelvir. M^pro^ is displayed as cartoon and colored in grey, mutant residues are shown as spheres and colored in blue. Nirmatrelvir is shown as ball and stick and colored in green. Selected residues are shown as lines and colored in cyan. Substrate binding subsites are labeled in red. (A) Mpro WT (PDBid 8DZ2). (B) Mpro Q189K (PDBid 8DZ6). (C) Mpro N142S (PDBid 8E26). (D) Mpro M49I (PDBid 8E25). (E) Mpro A193T (PDBid 8DZA). (F) Mpro A193S (PDBid 8E1Y). WT crystal structure is aligned with each M^pro^ mutant, displayed as grey ghost. Polar contacts are showed as black dashes.

In nirmatrelvir complexes, the S2 subsite is occupied by the dimethylcyclopropylproline substituent which interacts mainly by hydrophobic interactions with the main chains of R188 and the side chains of H41, M49, M165 and Q189 (Fig. 3). In the mutant M49I, we see no major structural impacts relative to the WT, as all the above-mentioned contacts are maintained. The structure of Q189K mutant in complex with nirmatrelvir showed that the replacement of Q189 with lysine implicated in the disruption of a salt bridge between the NE2 of glutamine and the O atom of the carbonyl group of nirmatrelvir, dislocating the ligand position compared to the WT crystal structure, and causing the loose of hydrogen contacts between pyrrolidone moiety and the O from amino acid F140 (Fig. 3).

The pyrrolidone group occupied the S1 pocket in both chains, except for mutant G143S where the pyrrolidone displays a higher flexibility, exhibiting four different binding modes for each chain (Fig. 4). The structure of mutant N142S did not reveal any significant structural differences, consistent with the similar enzymatic activity and drug inhibition observed for mutants N142S and N142D (Fig. 3).

**Figure 4.**
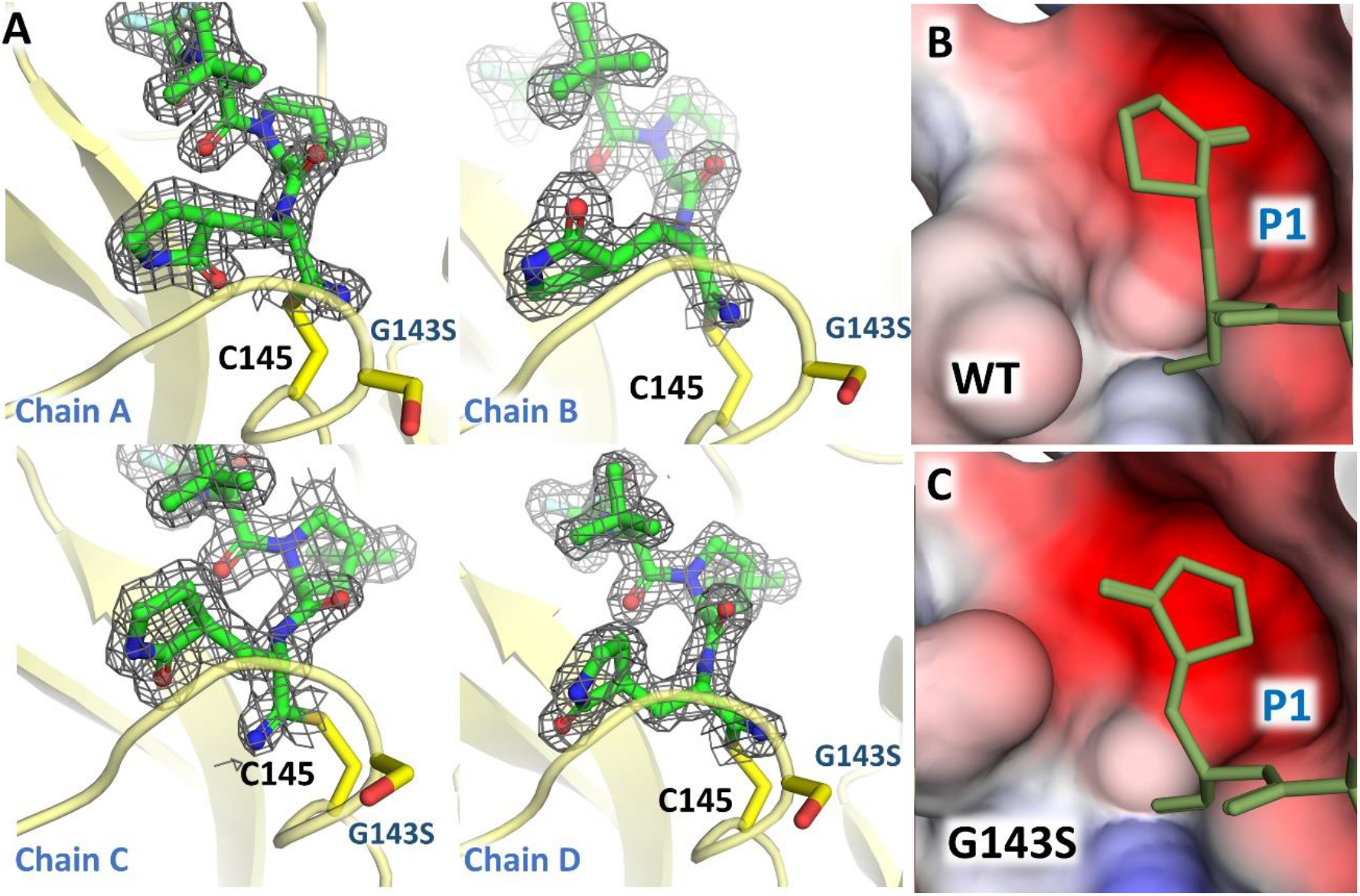
Crystal structure of G143S M^pro^ mutant in complex with nirmatrelvir. (A) Chains A-D of G143S M^pro^ in complex in Nirmatrelvir highlighting the different ligand conformations. M^pro^ is shown as yellow cartoon. Residues G143 and C145 are shown as sticks and colored in yellow. For Nirmatrelvir, the 2mFo-DFc electron density contoured at 1.0σ is represented in grey. (B) Surface charge representation of S1 subpocket for WT Mpro, evidencing the conformation of the pyrrolidine group. (C) Electrostatic charged calculated with APBS(41), projected on surface representation of S1 subpocket for G143S M^pro^ evidencing the conformation of the pyrrolidine group. For (B) and (C) Nirmatrelvir is shown as lines and colored in yellow.

The tert-butyl substituent at P3 position remained solvent exposed for all mutants, as observed for the WT crystal structure. The S4 sub-pocket accommodate the trifluoroacetamide substituent at position P4, where the main chain oxygen of the E166 residue makes a hydrogen bond with nirmatrelvir amide nitrogen N1 (Fig. 3). For both crystal structures with mutations of residue 193, A193S and A193T, even though being the most distant mutations of the active site, the additional steric volume and polar features of the serine and threonine side chains determined a displacement in the ligand binding mode in comparison with the binding mode to the WT. The binding mode of nirmatrelvir to both A193S and A193T mutants were similar, but the displacement is more severe in A193S, causing the disruption of pyrrolidone hydrogen bound with E166 (Fig. 3).

C145 and G143 residues are responsible for stabilizing the oxyanion hole, where the backbone nitrogen interacts through hydrogen bonds (Fig. 3). The mutant G143S complex revealed that the substitution of a glycine residue with a serine changed the charge distribution around S1 sub-pocket, causing multiple conformations of the pyrrolidone moiety due to the rotation of this moiety (Fig. 4). As a crystallographic consequence, the multiple conformations of the pyrrolidone caused the break of one of the orthorombic two-fold symmetry axis, shifting this crystal to a monoclinic space group with similar cell relative to the above (Table S2).

For ensitrelvir complexes, both structures, WT and M49I exhibited a similar non-covalent binding mode maintaining the key interactions with residues T26 (3.5 Å), G143 (3.2 Å), S144 (3.2 Å), C145S (3.3 Å), H163 (3.1 Å) and E166 (3.0 Å) through productive hydrogen bonds (Fig. 5). A characteristic of ensitrelvir in complex with M^pro^ is the rotation of H41 relative to the apo/nirmatrelvir structure, forming a face-to-face π stack with the 3,4,5-trifluorobenzene moiety of Ensitrelvir (Fig 5).

**Figure 5.**
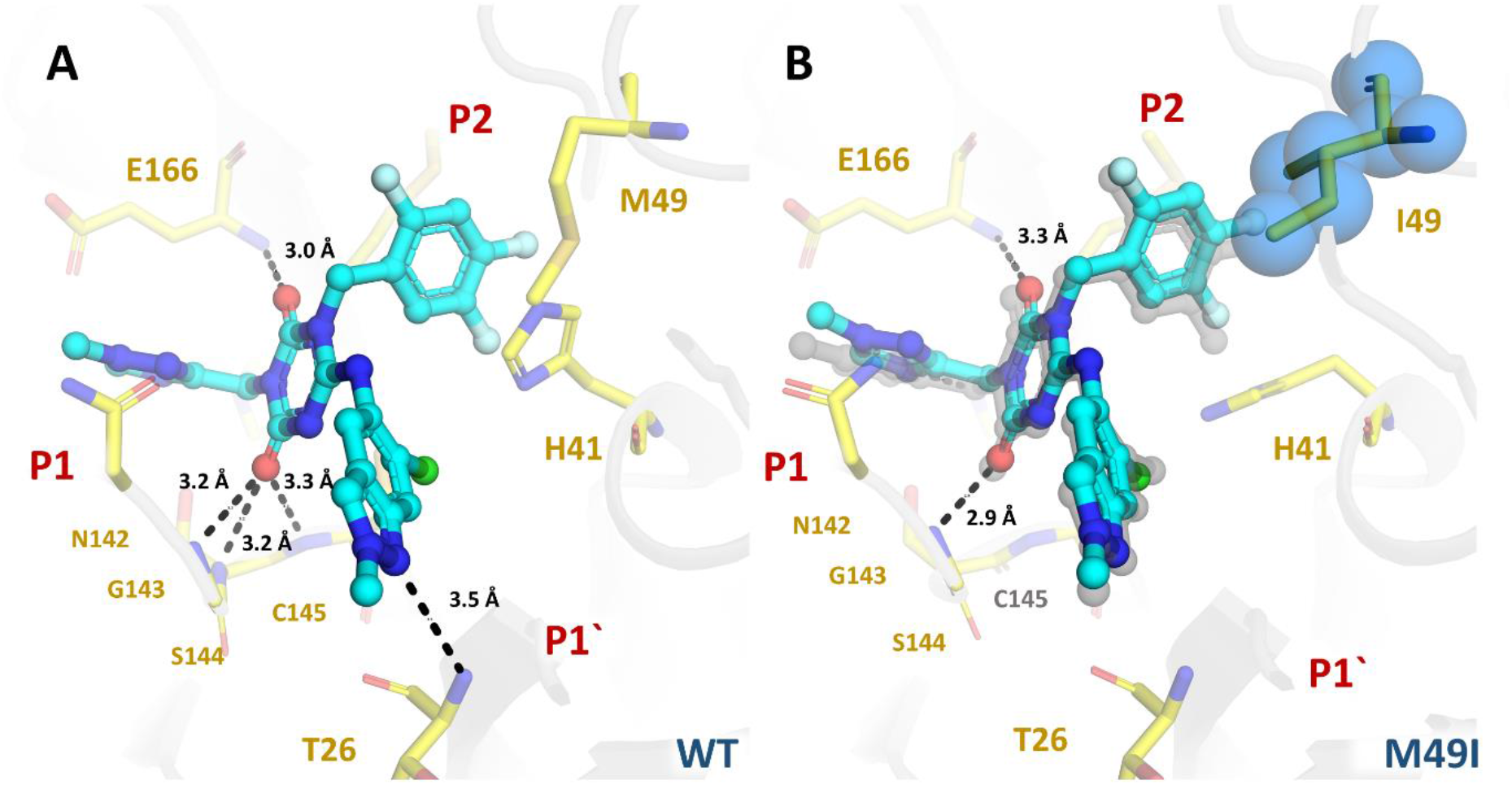
Crystal structures of M^pro^ in complex with ensitrelvir. M^pro^ is displayed as cartoon and colored in grey, mutant residue is shown as spheres and colored in blue. Selected residues are displayed as yellow sticks. Ensitrelvir is shown as ball and stick and colored in cyan. Substrate binding subsites are labeled in red. (A) Mpro WT (PDBid 8DZ0). (B) Mpro M49I (PDBid 8DZ1) aligned with M^pro^ WT, displayed as grey ghost. Polar contacts are showed as black dashes.

The crystal structure of M49I mutant revealed that the substitution dislocated the ligand orientation within the binding site towards the P2 position. The binding mode analysis suggested that this displacement is related to the greater hydrophobicity and steric volume of the isoleucine residue in comparison with the methionine residue. Additionally, for the WT structure, the side chain of the catalytic H41 residue is flipped undergoing a π-interaction with the trifluorophenyl substituent, whereas for the M49I mutant, the H41 rotamer is positioned similarly to M^pro^ apo structures and nirmatrelvir complexes (Fig. 5). The disruption of the ligand causes the break of hydrogen bonds between T26, S144 and C145 (Fig 5).

## Discussion

Due its critical role in viral replication and high conservation of catalytic residues, M^pro^ has proved to be a valuable target for the discovery and development of new antiviral candidates. Still, the high rate of infection of SARS-CoV-2 summed with the unprecedent global effort in sequencing and monitoring the virus evolution has revealed recurrent synapomorphies within a range of action of recently developed antivirals. Even though none of the characterized polymorphisms here are currently considered of interest by the World Health Organization due its low occurrence, the broad use of nirmatrelvir/ensitrelvir is likely to induce the selection of resistant viral strains. The kinetic characterization data collected herein showed that many of the selected polymorphisms resulted in enzymatically efficient forms of M^pro^, indicating that most of the observed mutations would be able to generate versions of M^pro^ with high fitness for selection (Fig. 2). Only G143S, Q189K and A193V mutants exhibited significantly decreased enzymatic efficiency against the fluorogenic substrate. In general, at least one of the inhibitors retained the inhibitory activity against most mutants, including the ones with higher occurrence (e.g., T190I, A191V, A193T and A193V) (Fig. 1 and 2B).

Nirmatrelvir is a covalent peptidomimetic inhibitor of SARS-CoV-2 M^pro^ developed by Pfizer, designed to compete with the substrate at the P1 and P2 subsites. To improve the inhibitory potency, the researchers had mimicked the glutamine P1 interactions using a pyrrolidinone moiety, interconnected by an amide bond with the 6,6-dimethyl-3-azabicyclo hexane moiety that mimics the hydrophobic interactions of P2 pocket (23). Moreover, the nitrile warhead is covalent bound to the catalytic Cys145 residue and makes hydrophobic interactions in P2. Nirmatrelvir key interactions with M^pro^ also include hydrogen bounds with H163, E166 and Q189. A recent study used deep mutational scanning to map the mutational landscape of M^pro^, and found that M49, N142, E166, P168, Q189 and A191 residues have high flexibility, and mutant E166V is highly resistant to nirmatrelvir (24). In another very recent pre-publication, the authors used a similar approach to ours to identify 11 mutants resistant to nirmatrelvir, including S144M/F/A/G/Y, M165T, E166Q, H172Q/F, and Q192T/S/V (25).

The mutant panel tested herein explored the capacity of nirmatrelvir to retain *in vitro* activity against circulating active polymorphisms of M^pro^. Our data indicated that nirmatrelvir showed nanomolar inhibitory activity against most of the tested mutants, including the most common mutation found, A191V, which showed a 2.6-fold increase in the *K*_i_ value for nirmatrelvir. The mutations that affect P2 subsite residues M49 and M165 had minor effect on nirmatrelvir activity (Fig 2B). The structure of M49I in complex with nirmatrelvir showed that the mutation has minor effect on ligand orientation with the binding site in comparison with the WT, where all key molecular contacts were maintained (Fig. 3D). More distant mutations affecting subsites P3-P4 small residues, such as A191T, A191V, A193T and A193S (but not A193V) have shown significantly increased *K*_i_ values for nirmatrelvir when compared with P1 and P2 mutations. The structure of A193T and A193S mutants showed that the increased steric volume and polar features of the side chains caused a displacement of the inhibitor related to the WT (Fig. 3E - F). As expected, we also notice that mutations in a specific amino acid seemed to have a greater effect in resistance than conserved mutations, as exemplified by R188K vs R188S (1.4 vs 2.9 -fold increase in *K*_i_). Another potentially impactful resistance mechanism depicted here was the one caused by the Q189K mutant, in which the crystal structure revealed that the absence of a productive hydrogen bond between the oxygen atom of the amide group of nirmatrelvir and the side chain of Q189 caused a rearrangement of the inhibitor’s binding mode (Fig. 3B), leading to an increase of 16-fold in the *K*_i_ value (Fig. 2B). Notwithstanding, it worth mentioning that the kinetic characterization indicated that this mutant showed 30 % of enzyme efficacy when compared with the WT (Fig. 2A), which should impact this polymorphism chance of being selected.

The most notorious structural differences observed in the mutant panel was for G143S mutant, in which the pyrrolidinone substituent of nirmatrelvir bound to M^pro^ in four distinct conformations on P1 (Fig. 4A). These variations seemed to break the orthorombic symmetry of the crystal system into monoclinic, adding one extra copy of M^pro^ dimer to the asymmetric unit (Fig. 4A). In this sense, we analyzed the charge distribution on P1 in the crystal structure of G143S in complex with nirmatrelvir and compare with the WT. Our findings indicated that the mutation had a broader distribution of the electron negative charges that forms the cavity at the WT (Fig. 4B). Thus, it is likely the mutation on G143 to serine shifted the charge distribution and/or stereochemistry on S1, preventing the stabilization of the pyrrolidinone group into a single conformation. A similar effect might explain why this mutation also depleted the enzyme efficacy versus a substrate containing Gln sidechain at P1. Despite the high increase in resistance generated by this mutation, the low catalytic efficiency exhibited by this polymorphism might limit its ability to be selected over other strains, unless this is associated with secondary mutations that compensate the deleterious effect of the mutation.

Ensitrelvir is nonpeptidic noncovalent inhibitor of SARS-CoV-2 M^pro^, developed by Shionogi using an intense structure-based drug design program (13). Their clinical candidate uses a methyl-triazole substituent (P1) linked to a trifluorophenyl group (P2) by a triazinane-2,4-dione, that also connects a third substituent containing a 6-chloro-2-methylindazol-5-amine that competes with P1’ subsite (13). This scaffold allowed ensitrelvir to achieve high enzymatic/antiviral activity and great metabolic stability, with key hydrogen bounds formed with T26, G143, H163 and E166 (13). In contrast with nirmatrelvir, ensitrelvir seems to be more susceptible to resistance in mutations affecting P2 subsite, such as M49I and M49T (Table 1). The structure of M49I in complex with ensitrelvir revealed that the higher hydrophobicity of isoleucine side chain caused a displacement of the inhibitor towards the P2 cavity, likely affecting its inhibitory activity (Fig 5B). Another key difference between ensitrelvir and nirmatrelvir resistance profile is that the former seems to retain near full activity against Q189K mutant.

The combined results allow us to conclude that these two distinct inhibitors have a different resistance profile against a panel of mutants, which can be explained by the distinct binding modes to M^pro^. These results are not only important in the monitoring of emergence of resistant strains of SARS-CoV-2, but also for planning a more suitable treatment in the event of one of these polymorphisms became a strain of concern. Moreover, the depicted complexes between inhibitors and mutants help us to understand the structural features involved in resistance, which should assist the development of the next generation of M^pro^ inhibitors.

## Methods

### 1. Identification of M^pro^ polymorphisms

To predict compound resistance, we selected polymorphic versions of M^pro^ that have been already identified in circulation. For that, we sort sequencing data available from GISAID hCoV-19/SARS-CoV-2 sequences database(22) in CoV-GLUE (http://cov-glue.cvr.gla.ac.uk) relative to M^pro^. Then, we selected all mutants that are within 7.0 Å radius of the active site with an n ≥ 10 individuals.

### 2. Site-directed mutagenesis, protein expression and purification

Cloning and expression were performed as described in Noske et al, 2021(7). The viral cDNA template (GenBank MT126808.1), kindly provided by Dr. Edison Durigon (University of São Paulo, São Paulo, Brazil), was synthetized using the SCRIPT One-Step RT-PCR kit (Cellco Biotec) and random hexamers primers. M^pro^ coding region (residues 3264-3569) was inserted into pET_M11/LIC vector using the LIC method(26). Final construct contains a N-terminal 6xHis-tag followed by a TEV site and the native N-terminal M^pro^ residues (YFQGAMSAVLQ↓SGFRK). For the site-directed mutagenesis, pET_M11/LIC-Mpro vector was used as template for inverse-PCR(27). All the PCRs were performed using FastPol polymerase (Cellco biotec) and primers from Table S1. PCR product was digested with DPNI (NEB), followed by treatment with T4 Polynucleotide kinase (PNK) (Thermo Fisher Scientific) and T4 DNA ligase (Cellco biotec). Mutations were confirmed by sequencing.

Herein, we used a self-cleavable construct of SARS-CoV-2 M^pro^ that was successfully expressed and purified using ammonium sulfate precipitation, followed by anion-exchange chromatography, with a final yield of 2.5 mg·L^−1^ of culture(7). The same expression and purification protocols were used to obtain M^pro^ mutants. All mutants were obtained with similar elution profile and final yield, except for N142L, that did not show an expression level suitable for purification. For protein expression, *E. coli* BL21 cells containing the recombinant plasmids were grown in ZYM 5052 medium(28) until OD_600_ reaches 0.6. Protein expression was induced by reducing the temperature to 18°C and cells were grown for 16h. Cells were harvested by centrifugation at 5000 x g, 4°C for 40 min and resuspended in lysis buffer (20 mM Tris pH 7.8, 150 mM NaCl, 1 mM DTT). Cells were disrupted by sonication. Lysate was clarified by centrifugation at 15.000 x g, 4°C for 30 min. After expression, M^pro^ is obtained in the native form after autocleavage. Protein was obtained by precipitation with addition of 1.5 M ammonium sulfate followed by incubation on ice for 10 min. Precipitated protein was isolated by centrifugation at 15.000 x g at 4 °C for 15 min. Protein was resuspended in lyses buffer and inject into a Superdex 200 26/100 (GE Healthcare) pre-equilibrated with gel filtration buffer (20 mM Tris pH 7.8, 50 mM NaCl, 1 mM DTT). After size exclusion chromatography, protein was buffer exchanged to 20 mM Tris pH 8.0, 1 mM DTT and further purified by ionic exchange chromatography using a Mono-Q column (GE Healthcare). Protein was eluted with a linear gradient of a buffer containing 20 mM Tris pH 8.0, 1 M NaCl and 1 mM DTT. Fractions containing the purified protein were collected and quantified using the measured absorbances at 280 nm and the theorical extinction coefficient of 32,890 M^−1^.cm^−1^. Protein purity was analyzed by SDS-PAGE. For enzymatic assays, protein was aliquoted at 0.5 mg/mL and flash-frozen using liquid nitrogen. Samples were stored at - 80° until use.

### 3. Protein crystallization and soaking

SARS-CoV-2 M^pro^ mutants were crystallized using sitting-drop vapor diffusion method. 1 µL of protein at 5-14 mg/mL was mixed with 1 µL of precipitating solution containing 0.1 M MES pH 6.7, 8% PEG 4K and 5% DMSO(29) and 0.2 µL of seed stock. Seed stock was obtained from orthorhombic crystal system M^pro^ crystals(7). Crystals were observed after 1-2 days at 16°C.

Crystals of SARS-CoV-2 M^pro^ mutants in complex with Nirmatrelvir and Ensitrelvir were obtained by soaking compound into M^pro^ mutants apo crystals grown as described previously. 2 µL of a solution containing 80% PEG 400, 20% DMSO and 10 mM of compound was added directly into the 2 µL crystallization drops. Crystallization plates were incubated at 16 °C for 24h. Crystals were manually harvested, and flash cooled in liquid nitrogen for data collection.

### 4. Data collection, processing, structure solving and refinement

Diffraction data collection was performed at MANACA beamline at the Brazilian Synchrotron SIRIUS (Campinas, Brazil) using a Pilatus 2M detector (Dectris). Data was processed and scaled using autoPROC/STARANISO from Global Phasing(30, 31). Resolution cut-off was determined by CC1/2(32, 33). For structure determination DIMPLE(34) was used for automated molecular replacement and initial refinement, using as initial template and search model the orthorhombic crystal structure of wild type M^pro^ (PDBid 7MBG). Ligand and covalent link restraints were generated using AceDRG through CCP4i2 program suite(35, 36). Refinement was conducted using REFMAC5(37) or Phenix.refine and manual rebuilding was performed using Coot(38). Structure validation was conducted using MolProbity(39). Figures were generated using Pymol (Schrödinger, LLC).

### 5. Activity and inhibition assays

Ensitrelvir was purchased from TCG lifesciences, whereas Nirmatrelvir was gently donated by Prof. Carlos A. Montanari. All enzymatic assays were performed using FRET-based substrate DABCYL-KTSAVLQ↓SGFRKM-E(EDANS)-NH2 in assay buffer (20 mM Tris pH 7.3, 1 mM EDTA, 1 mM DTT) in Corning^®^ 384-well white microplates. M^pro^ mutants were diluted to a final concentration of 40 nM. To determine the kinetics parameters (*K*_*m*_, V_max_ and k_cat_), the substrate was diluted to a range of concentrations from 200 μM to 0.1 μM. Reactions were previously incubated at 37°C for 10 min and started by addition of substrate in the respective concentrations. Fluorescence measures were monitored in SpectraMax Gemini EM Microplate Reader with λ_exc_/λ_emi_ of 360/460 nm, every 60 s over 60 min at 37º C. Initial velocity was derived from the slope of linear phase of each time-curse reaction, and Michaelis-Menten fitting was obtained using Origin Pro 9.5.1 Software (OriginLab). Relative efficiency of M^pro^ mutants was calculated by comparing the *K*_*m*_/k_cat_ relative to WT M^pro^. For IC_50_ determination, reactions containing Nirmatrelvir or Ensitrelvir from 10 µM to 0.0006 nM were previously incubated at 37°C for 10 min and started by addition of 10 µM of substrate. Inhibition percentages were determined by comparison with the DMSO control. All assays were performed in triplicates. For determining the *K*_i_ values, nirmatrelvir was considered an uncompetitive inhibitor, whereas ensitrelvir was considered a competitive inhibitor, and values were calculated using IC_50_-to-*K*_i_ (40).

## Accession numbers

Structure factors and atomic coordinates have been deposited with the protein data bank with accession codes PDB IDs 8DZ2, 8E25, 8E26, 8DZ9, 8DZ6, 8E1Y, 8DZA, 8DZ0 and 8DZ1. Other data are available from the corresponding author upon reasonable request.

## Acknowledgments

This research used facilities of the SIRIUS, part of the Brazilian Center for Research in Energy and Materials (CNPEM), a private non-profit organization under the supervision of the Brazilian Ministry for Science, Technology, and Innovations (MCTI). The MANACA beamline staff is acknowledged for the assistance during the experiments of proposal 20220605 and 20220741. Authors acknowledge Prof. Carlos Alberto Montanari for gently supplying nirmatrelvir. Funding: This project was funded by Coordenação de Aperfeiçoamento de Pessoal de Nível Superior (CAPES – Project 88887.516153/2020-00) and Fundação de Amparo à Pesquisa do Estado de São Paulo (FAPESP grants 2013/07600-3, 2015/16811-3, and 2016/19712-9). P.S. acknowledges also funding received from the Federal Ministry of Education and Research (BMBF) through KfW (Germany).

## Author contributions

ASG conceptualization, data analysis, writing. GDN, ESS, ID, MOG experimental, data analysis. RSF, GO, RVCG, PS data analysis, reviewing, funding.

## References

1. P. Zhou, et al., A pneumonia outbreak associated with a new coronavirus of probable bat origin. Nature 579, 270–273 (2020).

2. F. Wu, et al., A new coronavirus associated with human respiratory disease in China. Nature 579, 265–269 (2020).

3. Z. Jin, et al., Structure of Mpro from SARS-CoV-2 and discovery of its inhibitors. Nature 582, 289–293 (2020).

4. Y. Finkel, et al., The coding capacity of SARS-CoV-2. Nature 2020 589:7840 589, 125–130 (2020).

5. J. Lee, et al., Crystallographic structure of wild-type SARS-CoV-2 main protease acyl-enzyme intermediate with physiological C-terminal autoprocessing site. Nature Communications 2020 11:1 11, 1–9 (2020).

6. Z. Jin, et al., Structural basis for the inhibition of SARS-CoV-2 main protease by antineoplastic drug carmofur. Nat Struct Mol Biol 27, 529–532 (2020).

7. G. D. Noske, et al., A Crystallographic Snapshot of SARS-CoV-2 Main Protease Maturation Process. J Mol Biol, 167118 (2021).

8. W. Rut, et al., SARS-CoV-2 Mpro inhibitors and activity-based probes for patient-sample imaging. Nature Chemical Biology 2020 17:2 17, 222–228 (2020).

9. D. R. Owen, et al., An oral SARS-CoV-2 Mpro inhibitor clinical candidate for the treatment of COVID-19. Science, 374, 1586–1593 (2021).

10. L. Fu, et al., Both Boceprevir and GC376 efficaciously inhibit SARS-CoV-2 by targeting its main protease. Nat Commun 11, 1–8 (2020).

11. C. Ma, et al., Boceprevir, GC-376, and calpain inhibitors II, XII inhibit SARS-CoV-2 viral replication by targeting the viral main protease. Cell Res 30, 678–692 (2020).

12. J. D. A. Tyndall, S-217622, a 3CL Protease Inhibitor and Clinical Candidate for SARS-CoV-2. J Med Chem 65, 9, 6496–6498 (2022).

13. Y. Unoh, et al., Discovery of S-217622, a Noncovalent Oral SARS-CoV-2 3CL Protease Inhibitor Clinical Candidate for Treating COVID-19. J Med Chem 65, 6499–6512 (2022).

14. T. C. M. Consortium, et al., Open Science Discovery of Oral Non-Covalent SARS-CoV-2 Main Protease Inhibitor Therapeutics. bioRxiv, 2020.10.29.339317 (2021).

15. A. Narayanan, et al., Identification of SARS-CoV-2 inhibitors targeting Mpro and PLpro using in-cell-protease assay. Communications Biology 5:1 5, 1–17 (2022).

16. W. T. Harvey, et al., SARS-CoV-2 variants, spike mutations and immune escape. Nature Reviews Microbiology 2021 19:7 19, 409–424 (2021).

17. D. D. Singh, A. Parveen, D. K. Yadav, SARS-CoV-2: Emergence of New Variants and Effectiveness of Vaccines. Front Cell Infect Microbiol 11 (2021).

18. P. Mistry, et al., SARS-CoV-2 Variants, Vaccines, and Host Immunity. Front Immunol 12 (2022).

19. S. Ullrich, K. B. Ekanayake, G. Otting, C. Nitsche, Main protease mutants of SARS-CoV-2 variants remain susceptible to nirmatrelvir (PF-07321332) https:/doi.org/10.1101/2021.11.28.470226.

20. N. Krishnamoorthy, K. Fakhro, Identification of mutation resistance coldspots for targeting the SARS-CoV2 main protease. IUBMB Life 73, 670–675 (2021).

21. S. E. Greasley, et al., Structural basis for the in vitro efficacy of nirmatrelvir against SARS-CoV-2 variants. Journal of Biological Chemistry 298, 101972 (2022).

22. S. Khare, et al., GISAID’s Role in Pandemic Response. China CDC Weekly, 2021, Vol. 3, Issue 49, Pages: 1049-1051 3, 1049–1051 (2021).

23. D. R. Owen, et al., An oral SARS-CoV-2 Mpro inhibitor clinical candidate for the treatment of COVID-19. Science (1979) (2021) https:/doi.org/10.1126/SCIENCE.ABL4784 (November 7, 2021).

24. S. Iketani, et al., Functional map of SARS-CoV-2 3CL protease reveals tolerant and immutable sites. Cell Host Microbe (2022) https:/doi.org/10.1016/J.CHOM.2022.08.003 (August 15, 2022).

25. Y. Hu, et al., Naturally occurring mutations of SARS-CoV-2 main protease confer drug resistance to nirmatrelvir. bioRxiv, 2022.06.28.497978 (2022).

26. C. Aslanidis, P. J. De Jong, Ligation-independent cloning of PCR products (LIC-PCR). Nucleic Acids Res 18, 6069–6074 (1990).

27. W. Salaemae, M. Junaid, C. Angsuthanasombat, G. Katzenmeier, Structureguided mutagenesis of active site residues in the dengue virus two-component protease NS2B-NS3. 1–8 (2010).

28. F. W. Studier, Protein production by auto-induction in high density shaking cultures. Protein Expr Purif 41, 207–234 (2005).

29. A. Douangamath, et al., Crystallographic and electrophilic fragment screening of the SARS-CoV-2 main protease. Nat Commun 11 (2020).

30. C. Vonrhein, et al., Data processing and analysis with the autoPROC toolbox. Acta Crystallogr D Biol Crystallogr 67, 293–302 (2011).

31. C. Vonrhein, et al., Advances in automated data analysis and processing within autoPROC, combined with improved characterisation, mitigation and visualisation of the anisotropy of diffraction limits using STARANISO. Acta Crystallogr A Found Adv 74, a360–a360 (2018).

32. P. A. Karplus, K. Diederichs, Assessing and maximizing data quality in macromolecular crystallography. Curr Opin Struct Biol 34, 60–68 (2015).

33. P. A. Karplus, K. Diederichs, Linking crystallographic model and data quality. Science (1979) 336, 1030–1033 (2012).

34. M. D. Winn, et al., Overview of the CCP4 suite and current developments. urn:issn:0907-4449 67, 235–242 (2011).

35. F. Long, et al., AceDRG: A stereochemical description generator for ligands. Acta Crystallogr D Struct Biol 73, 112–122 (2017).

36. L. Potterton, et al., CCP4i2: the new graphical user interface to the CCP4 program suite. Acta Crystallogr D Struct Biol 74, 68 (2018).

37. G. N. Murshudov, et al., REFMAC5 for the refinement of macromolecular crystal structures. Acta Crystallogr D Biol Crystallogr 67, 355 (2011).

38. P. Emsley, B. Lohkamp, W. G. Scott, K. Cowtan, Features and development of Coot. Acta Crystallogr D Biol Crystallogr 66, 486–501 (2010).

39. V. B. Chen, et al., MolProbity: all-atom structure validation for macromolecular crystallography. Acta Crystallogr D Biol Crystallogr 66, 12–21 (2010).

40. R. Z. Cer, U. Mudunuri, R. Stephens, F. J. Lebeda, IC50-to-Ki: a web-based tool for converting IC50 to Ki values for inhibitors of enzyme activity and ligand binding. Nucleic Acids Res 37, W441–W445 (2009).

41. E. Jurrus, et al., Improvements to the APBS biomolecular solvation software suite. Protein Science 27, 112–128 (2018).

