## Supplemental material 1 for "Structural basis of nirmatrelvir and ensitrelvir resistance profiles against SARS-CoV-2 Main Protease naturally occurring polymorphisms"

Table S1. Primers used in inverse-PCR for each M^pro^ mutants.

| Polymorphism | Forward | Reverse |
| --- | --- | --- |
| M49I | CCTCTGAAGACATTCTTAACCC | TGCAGATCACATGTCTTGG |
| M49T | CTCTGAAGACACGCTTAACCC | GTGCAGATCACATGTCTTGGA |
| N142S | CATTCCTTAGTGGTTCATGTGG | AACCCTTAATAGTGAAATTGGGC |
| N142D | CATTCCTTGATGGTTCATGTG | AACCCTTAATAGTGAAATTGGGC |
| N142L | CATTCCTTCTTGGTTCATGTG | AACCCTTAATAGTGAAATTGGGC |
| G143S | CATTCCTTAATAGTTCATGTGGTAGTG | AACCCTTAATAGTGAAATTGGGC |
| M165I | CATGCACCATATTGAATTACCAAC | TAACAAAAAGAGACACAGTCATAATCTATGTT |
| R188K | GTTGACAAGCAAACAGCAC | AAAAGGTCCATAAAAGTTACCTTC |
| R188S | GTTGACAGCCAAACAGC | AAAAGGTCCATAAAAGTTACCTT |
| Q189K | GACAGGAAAACAGCACAAG | AACAAAAGGTCCATAAAAGTTAC |
| T190I | GGCAAATTGCACAAGC | TGTCAACAAAAGGTCCATAAA |
| A191T | GCAAACAACACAAGCAGCTG | CTGTCAACAAAAGGTCCATAAAAGTTAC |
| A191V | GGCAAACAGTACAAGCAGCTG | TGTCAACAAAAGGTCCATAAAAGTTAC |
| A193S | GCACAATCAGCTGGTACG | TGTTTGCCTGTCAACAAAA |
| A193T | GCACAAACAGCTGGTACG | TGTTTGCCTGTCAACAAAAG |
| A193V | CAGCACAAGTAGCTGGTACGG | TTTGCCTGTCAACAAAAGGTC |

Table S2. Data collection and refinement statistics for Mpro mutants in complex with Nirmatrelvir. Statistics for the highest-resolution shell are shown in parentheses.

|  | | WT | M49I | N142S | G143S | Q189K | A193S | A193T |
| --- | --- | --- | --- | --- | --- | --- | --- | --- |
| *Data collection* | | MANACA beamline | MANACA beamline | MANACA beamline | MANACA beamline | MANACA beamline | MANACA beamline | MANACA beamline |
| Space group | | *P*2_1_2_1_2_1_ | *P*2_1_2_1_2_1_ | *P*2_1_2_1_2_1_ | *P2_1_* | *P*2_1_2_1_2_1_ | *P*2_1_2_1_2_1_ | *P*2_1_2_1_2_1_ |
| Cell dimensions | |  |  |  |  |  |  |  |
|  | *a,b, c* (Å) | 67.319, 97.536, 102.327 | 67.859, 100.068, 103.533 | 67.841, 100.619, 103.356 | 67.686, 102.808, 101.554 | 67.782, 101.572, 103.561 | 68.025, 100.905, 103.025 | 67.989, 99.979, 103.638 |
|  | *α, β, γ* (^o^) | 90, 90, 90 | 90, 90, 90 | 90, 90, 90 | 90, 91.2, 90 | 90, 90, 90 | 90, 90, 90 | 90, 90, 90 |
| Resolution range (Å) | | 70.6 - 2.13 (2.20 - 2.13) | 72.06 - 1.87 (1.937 - 1.87) | 56.31-1.84  (1.91-1.84) | 101.53-1.66 (1.72-1.66) | 72.52-2.37  (2.45-2.37) | 72.43-2.48  (2.57-2.48) | 72.06-1.96  (2.03-1.96) |
| Unique Reflections | | 37960 (3779) | 30317 (138) | 24397 (7) | 74929 (112) | 16996 (78) | 15497 (86) | 29962 (85) |
| Multiplicity | | 5.1 (5.2) | 5.6 (5.9) | 4.0 (3.3) | 3.2 (2.9) | 5.3 (4.3) | 6.4 (3.9) | 12.9 (11.0) |
| Completeness (%) | | 98.60 (99.58) | 51.42 (2.37) | 39.62 (0.1) | 46.1 (0.7) | 56.9 (2.7) | 59.5 (3.4) | 58.3 (1.7) |
| *I*/σ*I* | | 8.6 (2.3) | 7.4 (1.6) | 4.4 (1.5) | 3.7 (1.7) | 4.6 (1.7) | 3.9 (1.3) | 7.8 (1.7) |
| *R*_p.i.m._ (%)^1^ | | 0.065 (0.365) | 0.084 (0.377) | 0.134 (0.250) | 0.121 (0337) | 0.109 (0.336) | 0.127 (0.467) | 0.080 (0.469) |
| CC_1/2_^2^ | | 0.991 (0.763) | 0.989 (0.684) | 0.949 (0.823) | 0.968 (0.463) | 0.997 (0.683) | 0.965 (0.575) | 0.991 (0.253) |
| *Refinement* | |  |  |  |  |  |  |  |
| *R*_work_/*R*_free_ ^3^ | | 0.19/0.23 | 0.20/0.25 | 0.23/0.27 | 0.21/0.25 | 0.22/0.30 | 0.22/0.27 | 0.20/0.24 |
| Number of atoms | |  |  |  |  |  |  |  |
|  | Waters | 259 | 202 | 219 | 640 | 74 | 31 | 175 |
|  | Ligands | 82 | 82 | 78 | 144 | 82 | 78 | 90 |
|  | Protein residues | 4691 | 4690 | 4742 | 9366 | 4687 | 4674 | 4691 |
| RMS(bonds) (Å) | | 0.013 | 0.014 | 0.0126 | 0.0153 | 0.0129 | 0.0128 | 0.0133 |
| RMS(angles) (^o^) | | 1.91 | 1.82 | 1.66 | 2.09 | 1.78 | 1.68 | 1.77 |
| Ramachandran favored (%) | | 96.68 | 95.99 | 97.70 | 94.42 | 94.20 | 95.99 | 97.01 |
| Ramachandran outliers (%) | | 0.33 | 0 | 0.33 | 0.33 | 0.33 | 0.33 | 0.17 |
| Clashscore^4^ | | 6.39 | 4.15 | 7.28 | 15.3 | 7.14 | 7.28 | 7.98 |
| Average *B*-factors (Å²) | | 37.08 | 35.91 | 19.02 | 20.77 | 36.54 | 33.23 | 36.48 |
|  | Macromolecules | 34.98 | 35.78 | 19.72 | 20.38 | 35.84 | 32.28 | 34.78 |
|  | Ligands | 42.37 | 48.17 | 24.47 | 19.99 | 52.7 | 65.78 | 48.84 |
|  | Solvent | 42.40 | 33.79 | 20.64 | 26.3 | 24.71 | 21.57 | 36.23 |
| PDB code | | 8DZ2 | 8E25 | 8E26 | 8DZ9 | 8DZ6 | 8E1Y | 8DZA |

**1** *Rp.i.m.* = Σhkl {1/[N(hkl) - 1]}1/2 x Σj│Ii(hkl) - <(hkl)> │/ Σhkl Σj Ii(hkl)4.

**2** CC1/2 is the correlation coefficient determined by two random half data sets5

**3** *Rwork* = Σhkl│Fo(hkl) - Fc(hkl)│/ Σhkl Fo(hkl). *Rfree* was calculated for a test set of reflections (10%) omitted from the refinement.

**4** Clashscore is the number of clashes calculated for the model per 1000 atoms**.**

Table S3. Data collection and refinement statistics for Mpro mutants in complex with Ensitrelvir. Statistics for the highest-resolution shell are shown in parentheses.

|  | | WT | M49I |
| --- | --- | --- | --- |
| *Data collection* | | MANACA beamline | MANACA beamline |
| Space group | | *P*2_1_2_1_2_1_ | *P*2_1_2_1_2_1_ |
| Cell dimensions | |  |  |
|  | *a,b, c* (Å) | 67.9, 99.0, 103.1 | 67.5, 99.0, 103.3 |
|  | *α, β, γ* (^o^) | 90, 90, 90 | 90, 90, 90 |
| Resolution range (Å) | | 71.4 – 2.28 (2.6 – 2.28) | 71.5 – 2.0 (2.3-2.0) |
| Unique Reflections | | 15346 (767) | 25100 (1256) |
| Multiplicity | | 5.7 (5.6) | 5.0 (4.5) |
| Completeness (%) | | 89.4 (65.6) | 90.1 (65.5) |
| *I*/σ*I* | | 4.3 (1.6) | 6.2 (1.6) |
| *R*_p.i.m._ (%)^1^ | | 0.125 (0.438) | 0.1 (0.53) |
| CC_1/2_^2^ | | 0.973 (0.7) | 0.98 (0.54) |
| *Refinement* | |  |  |
| *R*_work_/*R*_free_ ^3^ | | 0.22/0.27 | 0.22/0.25 |
| Number of atoms | |  |  |
|  | Waters | 44 | 82 |
|  | Ligands | 86 | 86 |
|  | Protein residues | 4742 | 4742 |
| RMS(bonds) (Å) | | 0.005 | 0.006 |
| RMS(angles) (^o^) | | 0.78 | 0.93 |
| Ramachandran favored (%) | | 91.9 | 94.7 |
| Ramachandran outliers (%) | | 1.16 | 0.5 |
| Clashscore^4^ | | 11.16 | 8.32 |
| Average *B*-factors (Å²) | |  |  |
|  | Macromolecules | 43.43 | 41.46 |
|  | Ligands | 61.66 | 75.8 |
|  | Solvent | 30.51 | 39.0 |
| PDB code | | 8DZ0 | 8DZ1 |

**1** *Rp.i.m.* = Σhkl {1/[N(hkl) - 1]}1/2 x Σj│Ii(hkl) - <(hkl)> │/ Σhkl Σj Ii(hkl)4.

**2** CC1/2 is the correlation coefficient determined by two random half data sets5

**3** *Rwork* = Σhkl│Fo(hkl) - Fc(hkl)│/ Σhkl Fo(hkl). *Rfree* was calculated for a test set of reflections (10%) omitted from the refinement.

**4** Clashscore is the number of clashes calculated for the model per 1000 atoms**.**

**Fig S1.** A and B show Michaelis-Menten plots of all mutants. C and D show IC_50_ plots determination of Nirmatrelvir against the mutant panel. E and F show IC_50_ plots determination of Ensitrelvir against the mutant panel.

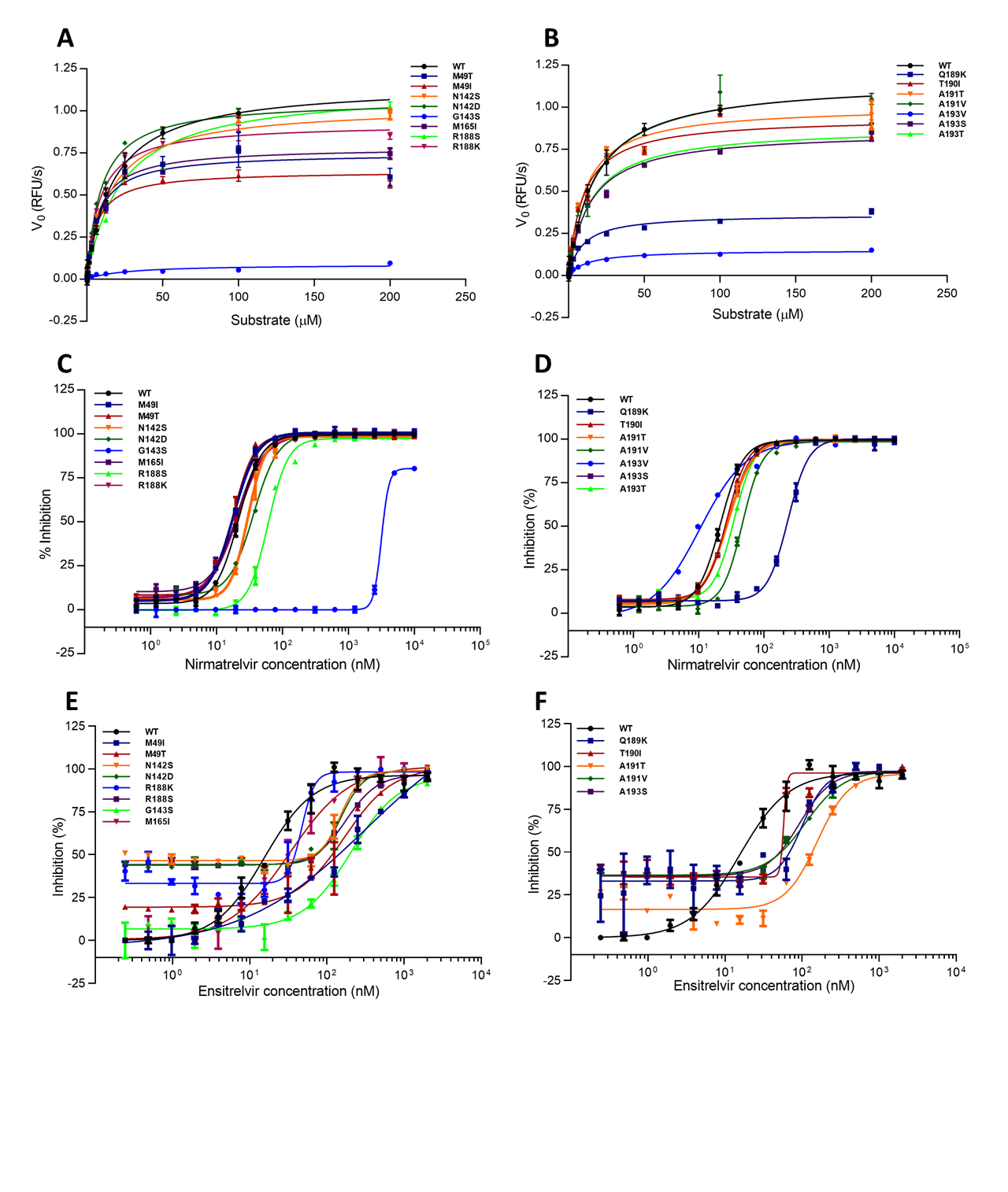
